# Avian influenza A viruses reassort and diversify differently in mallards and mammals

**DOI:** 10.1101/2021.02.08.430042

**Authors:** Ketaki Ganti, Anish Bagga, Juliana DaSilva, Samuel S. Shepard, John R. Barnes, Susan Shriner, Katia Koelle, Anice C. Lowen

## Abstract

Reassortment among co-infecting influenza A viruses (IAVs) is an important source of viral diversity and can facilitate expansion into novel host species. Indeed, reassortment played a key role in the evolution of the last three pandemic IAVs. Observed patterns of reassortment within a coinfected host are likely to be shaped by several factors, including viral load, the extent of viral mixing within the host and the stringency of selection. These factors in turn are expected to vary among the diverse host species that IAV infects. To investigate host differences in IAV reassortment, here we examined reassortment of two distinct avian IAV within their natural host (mallards) and a mammalian model system (guinea pigs). Animals were co-inoculated with A/wildbird/California/187718-36/2008 (H3N8) and A/mallard/Colorado/P66F1-5/2008 (H4N6) viruses. Longitudinal samples were collected from the cloaca of mallards or the nasal tract of guinea pigs and viral genetic exchange was monitored by genotyping clonal isolates from these samples. Relative to those in guinea pigs, viral populations in mallards showed higher frequencies of reassortant genotypes and were characterized by higher genotype richness and diversity. In line with these observations, analysis of pairwise segment combinations revealed lower linkage disequilibrium in mallards as compared to guinea pigs. No clear longitudinal patterns in richness, diversity or linkage disequilibrium were present in either host. Our results reveal mallards to be a highly permissive host for IAV reassortment and suggest that reduced viral mixing limits avian IAV reassortment in a mammalian host.

## Introduction

Influenza A viruses (IAVs) infect a broad range of host species. Many diverse lineages circulate in waterfowl (Anseriformes) and shorebirds (Charadriformes), with 16 hemagglutinin and 9 neuraminidase subtypes represented. This diverse viral gene pool is the ancestral source of IAV lineages circulating in poultry, swine, humans and other mammalian hosts (1, 2). Although host barriers to infection limit the range of hosts within which a given IAV lineage circulates, spillovers occur occasionally and can seed novel lineages (3, 4). When a novel IAV lineage is established in humans, the result is a pandemic of major public health consequence (5, 6).

The segmented nature of the IAV genome allows facile genetic exchange between viruses that co-infect the same host: through reassortment of intact gene segments, mixed infections frequently give rise to viral genotypes that differ from both parental strains (7). The potential for viral diversification through this mechanism is high. If co-infecting parental viruses differ in all eight gene segments, reassortment can yield 256 distinct viral genotypes from a single co-infected cell. However, the contribution of reassortment to IAV diversity can be limited by two major forms of constraint. The first is a lack of opportunity for reassortment between distinct strains. For reassortment to occur, viruses must infect the same host and the same cell within that host. These prerequisites will be met routinely only if viral spread is well-mixed and high density at the population and within-host levels, respectively. The second form of constraint is negative selection. Reassortment may be deleterious because it can break epistatic interactions among gene segments and the proteins they encode. As a result, progeny viruses with chimeric genotypes can be less fit than both parental strains (8, 9).

Despite these constraints, reassortment has repeatedly been implicated in the evolution of novel IAV lineages and is strongly associated with IAV host switching (10). Reassortment involving human seasonal strains and IAV adapted to avian and/or swine hosts led to the 1957, 1968, and most recently, 2009 influenza pandemics. The establishment in poultry of H7N9 and H5N1 subtype viruses that are highly pathogenic to humans followed from reassortment between enzootic poultry viruses and strains introduced transiently from wild birds (11–14). More recently, reassortment of poultry H5N1 viruses with IAV in wild birds led to the spread of highly pathogenic H5N8 and H5N2 subtype viruses to North America (15). Thus, reassortment can have major consequences for IAV evolution and host range expansion.

In the work described here, we used experimental IAV coinfection to evaluate the efficiency of reassortment in mallards, a major natural host of IAV. The viral strains used for coinfection, of H3N8 and H4N6 subtypes, are typical of viruses isolated from mallards and representative of lineages co-circulating in North American waterfowl. Both the hemagglutinin (HA) and neuraminidase (NA) subtypes represented are furthermore associated with frequent mixed infection within this ecological niche (16). We compared reassortment observed in this natural host-virus pairing to that seen with the same viral strains in a guinea pig model. In this way, the impact of host species on the extent of within-host genetic diversity achieved through reassortment was evaluated. Our results revealed abundant reassortment in mallards, giving rise to high genotype richness and diversity and low linkage disequilibrium between segments. In guinea pigs, reassortment rates were lower, fewer unique genotypes were detected and relatively high prevalence of parental genotypes resulted in low diversity. These findings indicate that mallards are highly permissive for IAV reassortment, whereas mammalian hosts may present a less permissive environment for IAV reassortment. Additional analyses suggest that lower levels of reassortment in guinea pigs stem from reduced viral mixing in this host, rather than pervasive purifying selection limiting the frequency of reassortants.

## Results

### Robust infection in co-inoculated mallards and guinea pigs

The genomes of the two viral strains used for co-infection, influenza A/wildbird/California/187718-36/2008 (H3N8) and A/mallard/Colorado/P66F1-5/2008 (H4N6) viruses, were sequenced in full and found to be genetically distinct in all eight segments (**Table 1**). The proteins encoded by these gene segments exhibited between 44.2% and 100% identity in their amino acid composition. The viral genome sequences were used to design eight segment-specific primer sets suitable for differentiation of the H3N8 and H4N6 gene segments by high resolution melt analysis.

**Table 1:**
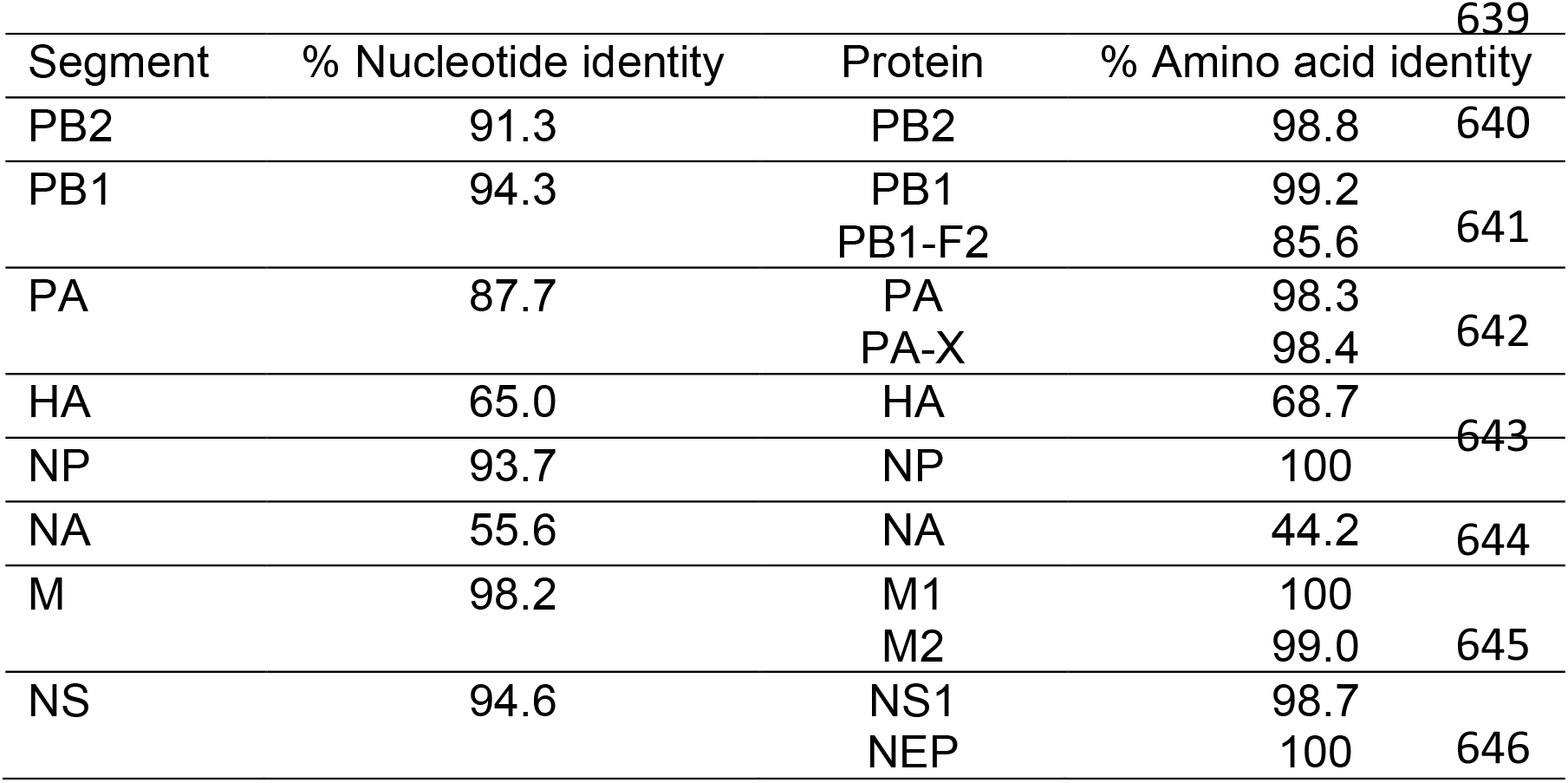
**Nucleotide and amino acid differences between A/wildbird/California/187718-36/2008 (H3N8) and A/mallard/Colorado/P66F1-5/2008 (H4N6) viruses**.

To evaluate the patterns of IAV reassortment *in vivo* and its reliance on host species, groups of eight mallards and eight guinea pigs were co-inoculated the H3N8 and H4N6 viruses and shed virus was sampled daily. Viral growth in each species was robust, but differed in the kinetics observed (**Figure 1**). The average peak titer reached in mallards and guinea pigs was comparable, at 4.5 and 5.2 log_10_PFU/mL, respectively, although guinea pigs showed a much broader range of values. Kinetics were more rapid in mallards, with peak viral load seen at the earliest time point (1 day post-inoculation) and clearance observed at 4 or 5 days post-inoculation. In guinea pigs, titers generally peaked at 2 days post-inoculation and shedding ceased at 5 days post-inoculation at the earliest.

**Figure 1:**
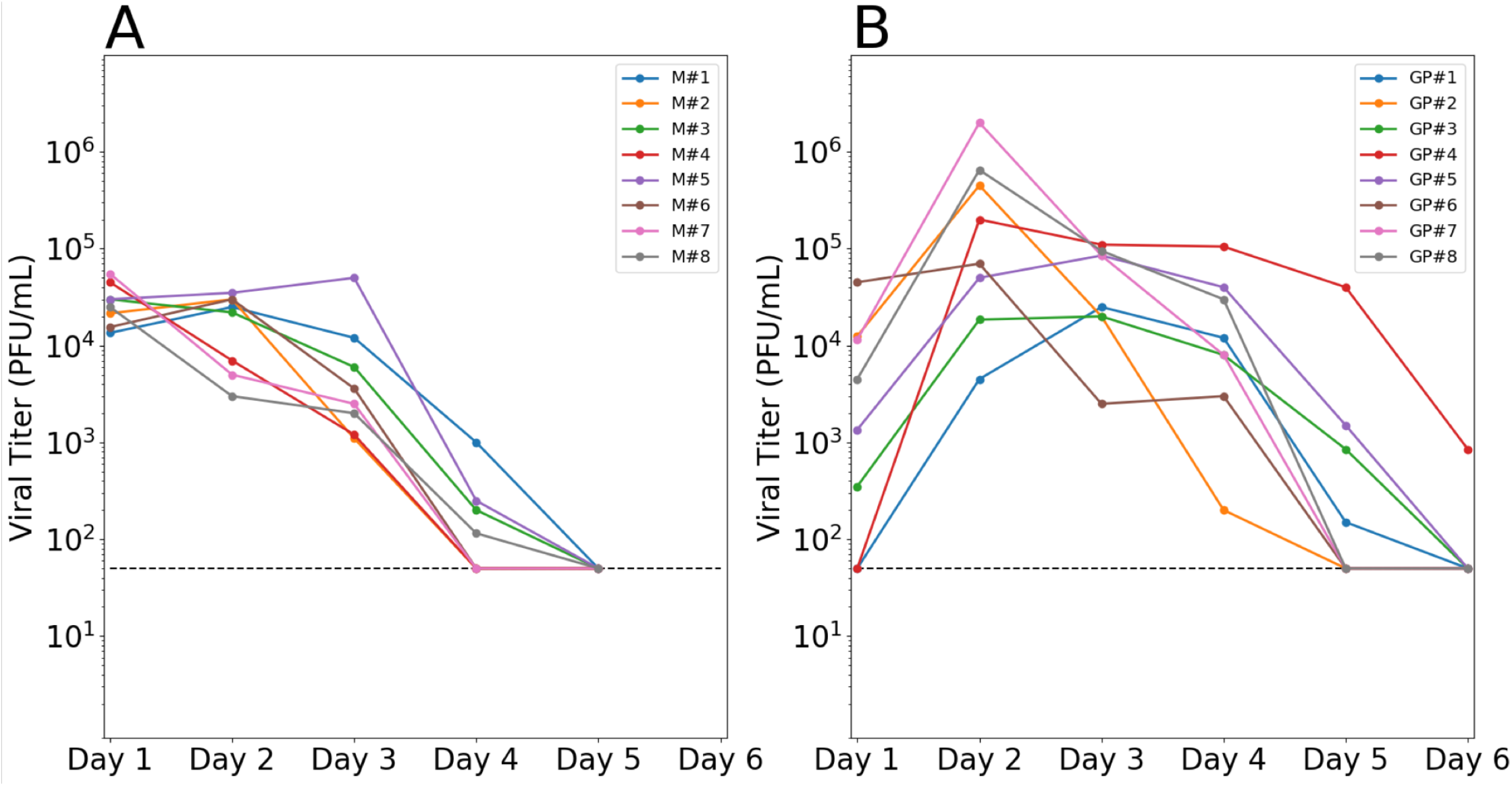
Efficient viral replication was observed in mallards and guinea pigs. Infectious viral titers in A) cloacal swab samples collected from mallards and B) nasal lavage samples collected from guinea pigs are plotted against day post-inoculation. Each colored line represents an individual animal. Horizontal dashed line shows the limit of detection.

### Genotypic diversity was higher in mallards than in guinea pigs

To quantify reassortment following coinfection, clonal isolates were derived from mallard cloacal swabs collected on days 1, 2 and 3 and from guinea pig nasal washes collected on days 2, 3 and 4. These time points were chosen because all individuals in the respective groups were shedding above the limit of detection on each of these days. Twenty-one clonal isolates were derived per sample. Genotyping was then performed for all eight gene segments of these isolates to evaluate whether they were of H3N8-origin or H4N6-origin. For a small minority of isolates, one or more gene segments gave an ambiguous result in our genotyping assay and the full isolate was therefore excluded from the analysis (giving a final sample size of 18-21 isolates per sample). In all individuals of both species, and at all time points examined, reassortment was detected (**Supplementary Figure 1**).

To characterize the outcomes of IAV coinfection in each host, the genotypes from the clonal isolates were quantitatively analyzed. Specifically, for each sample collected from a given individual on a given day, the frequency of unique genotypes, frequency of parental genotypes, genotype richness, and genotype diversity were determined (**Figure 2**). In mallards, the frequency of a given genotype was variable across time, reflecting infrequent carry-over of specific reassortant genotypes within a given individual (**Figure 2A** and **Supplementary Figure 2**). Parental genotypes were detected in each of the mallards on at least one sampling day, with frequencies on a given day varying between 0 and 0.6 (**Figure 2B**). Genotype richness, or the number of distinct genotypes detected in a sample, was typically high in mallards, with most samples showing richness values between 12 and the maximum value of 21 (**Figure 2C**). Diversity was measured using the Shannon-Weiner index, which considers both richness and evenness in the frequency with which genotypes are detected. Diversity in mallards tracked closely with richness and was also high, with all mallards showing diversity values greater than 2.3 on at least one sampling day (**Figure 2D**). Similar to mallards, genotype frequencies in guinea pigs were variable across time, again reflecting infrequent carry-over of specific reassortant genotypes (**Figure 2E** and **Supplementary Figure 2**). In striking contrast to mallards, however, parental genotypes were often predominant in guinea pigs, although the frequencies observed ranged widely, from 0 to 0.95 (**Figure 2F**). Genotype richness in guinea pigs was generally lower than that seen in mallards, with most samples comprising fewer than 12 unique genotypes (**Figure 2G**). Correspondingly, genotype diversity also appeared lower in guinea pigs than in mallards, with only one of the guinea pigs showing diversity above 2.3 and most values falling in the range of 0.5 to 2.0 (**Figure 2H**). Lower levels of genotype diversity in guinea pigs than in mallards were found to be statistically significant (p = 1.3×10^−7^; ANOVA).

**Figure 2:**
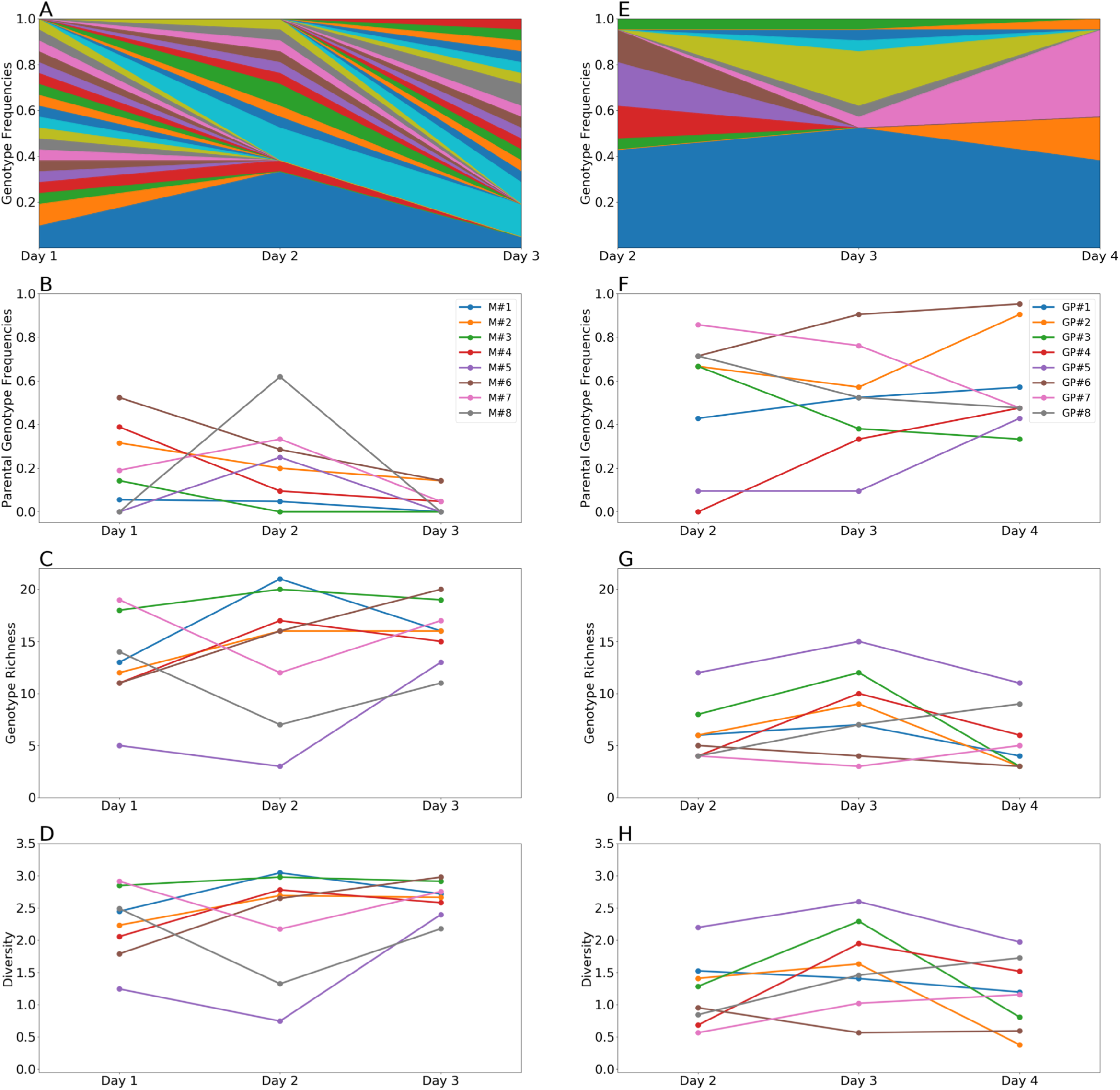
Viral populations in mallards showed lower frequencies of parental genotypes, higher genotype richness and higher diversity than those in guinea pigs. Results from mallards are shown in panels A-D and from guinea pigs in panels E-H. A, E) Stacked plot showing frequencies of unique genotypes detected in one representative animal over time. The lowermost two sections, in blue and orange, represent the H3N8 and H4N6 parental genotypes, respectively. B, F) Frequency of parental genotypes over time. The total frequency of H3N8 and H4N6 parental genotypes is plotted. C, G) Genotype richness over time. D, H) Diversity over time, as measured by the Shannon-Weiner index.

In the combined data set from both species, we noted an inverse correlation between diversity and parental genotype frequency (**Figure 3A**). To test the hypothesis that low diversity is driven by high parental frequencies, we therefore calculated viral evenness in each sample with and without the inclusion of parental genotypes. Evenness, rather than diversity, was analyzed in this way because the reduction in sample size brought about by excluding parental genotypes confounds effects on diversity. Since this exclusion typically has little impact on richness (reducing the number of unique genotypes by 0-2), any impact on diversity of removing parental genotypes would occur mainly through changes in evenness. Upon exclusion of parental genotypes, evenness in guinea pigs increased, while that in mallards was less affected (**Figure 3B, C**). Nevertheless, evenness was significantly lower in guinea pigs compared to mallards whether parental genotypes were included or not (p=1.47×10^−5^ and 0.0166, respectively, unpaired t-test). Thus, high parental genotype frequency contributes, but does not fully account for reduced viral diversity in guinea pigs compared to mallards.

**Figure 3:**
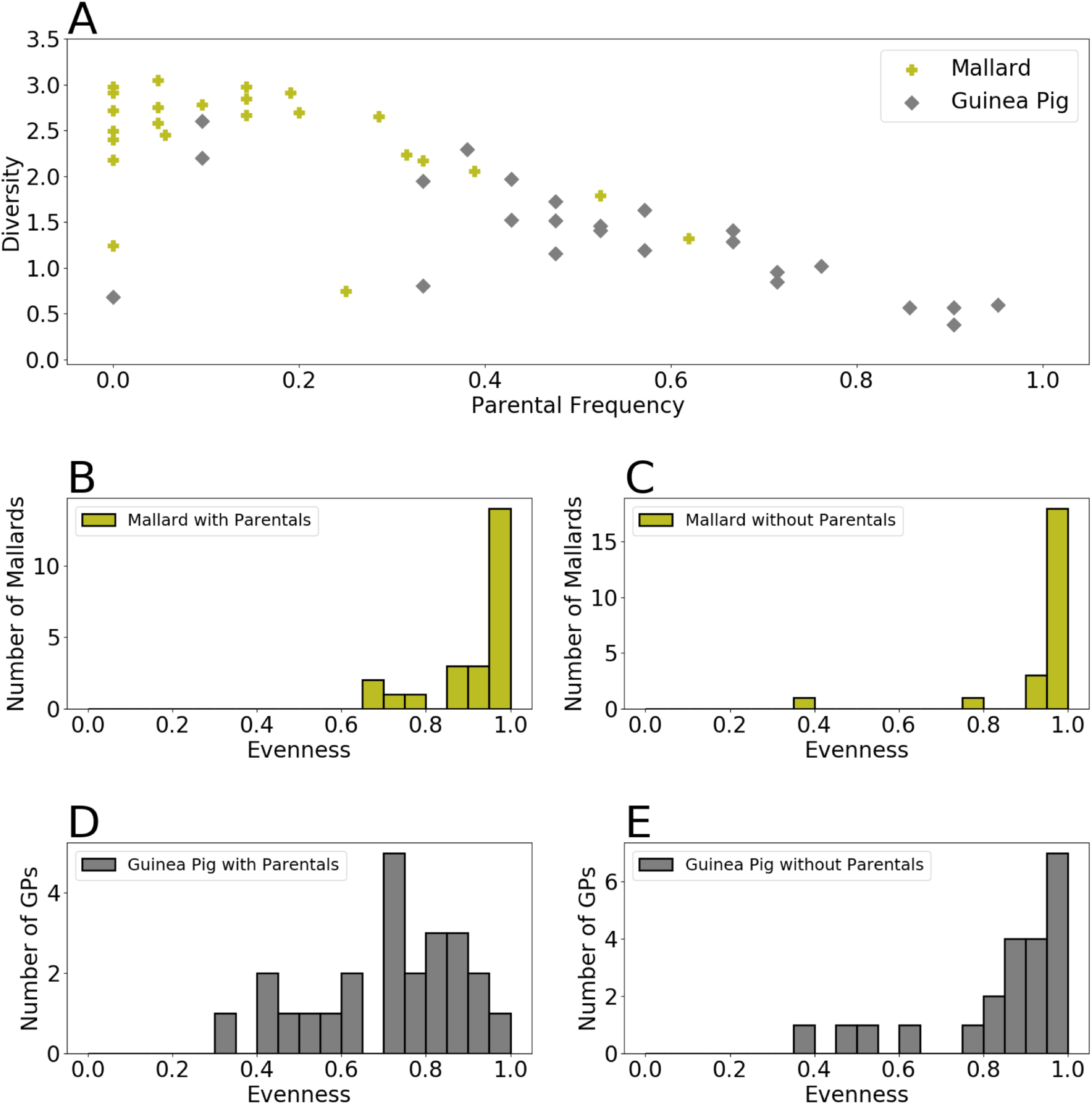
Lower levels of viral diversity in guinea pigs is only partially accounted for by high frequency of parental genotypes. A) Scatter plot of Shannon-Weiner diversity against parental genotype frequency. B) Histograms showing evenness of virus samples in mallards. C) Histograms showing evenness of virus samples in mallards, following removal of parental genotypes. D) Histograms showing evenness of virus samples in guinea pigs. E) Histograms showing evenness of virus samples in guinea pigs, following removal of parental genotypes.

### Parental genotypes drive higher linkage disequilibrium in guinea pigs than in mallards

To assess the extent to which reassortment was shaped by genetic linkages among segments, and whether this effect varied with host species, linkage disequilibrium (*D*) was evaluated for each pair of viral gene segments. This analysis examines the extent to which segments assorted non-randomly. For the pairing of PB2 and PB1, for example, four genotypes are possible: PB2_H3N8_PB1_H3N8_, PB2_H3N8_PB1_H4N6_, PB2_H4N6_PB1_H3N8_, and PB2_H4N6_PB1_H4N6_. If these four genotypes are present in a given viral population at the expected frequencies based on the prevalence of the four gene segments, *D* will equal zero, indicative of random assortment. Positive associations, in which specific genotypes are over-represented, will yield *D* > 0, while negative associations will give *D* < 0. In both mallards and guinea pigs, pairwise *D* values tended to be positive (**Figure 4**). Comparison between host species for each pairwise combination revealed that *D* was typically lower in mallards than guinea pigs (Mann Whitney U test, p<0.05 for 16 of the 28 segment pairings). While this result could suggest that reassortment in guinea pigs was more strongly shaped by positive segment associations than that in mallards, high linkage disequilibrium could alternatively be driven by a lack of opportunity for reassortment, leading to frequent occurrence of parental segment pairings.

**Figure 4:**
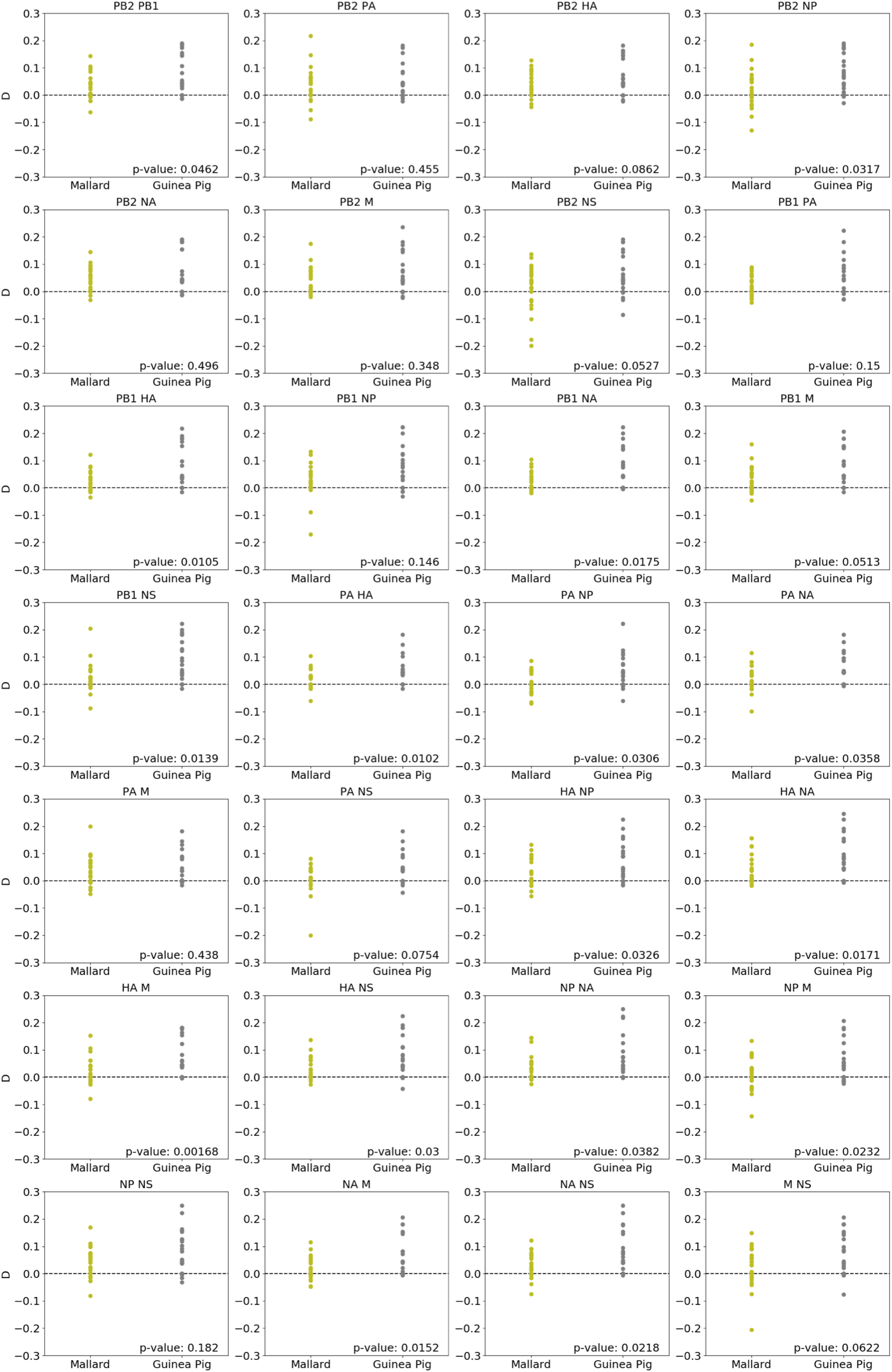
Pairwise linkage disequilibrium was typically lower in mallards than in guinea pigs. Linkage disequilibrium for each pairwise combination of IAV gene segments is shown. Each data point represents a mallard or guinea pig, with samples from all three time points aggregated for a given individual. P values indicate results of a Mann Whitney U test.

### Genotype patterns are consistent with random sampling of gene segments

To gauge whether the lower levels of reassortment in guinea pigs compared to ducks is due to a lack of reassortment opportunity versus more stringent selection in guinea pigs, we assessed the types of reassortants that were observed in guinea pigs and in mallards. Reasoning that less chimeric genotypes, in which only one or two segments are derived from a second parent strain, would be less likely to suffer fitness defects due to epistasis, we tested the hypothesis that such reassortants were over-represented in our data sets relative to others, and that they would be particularly overrepresented in guinea pigs relative to mallards. The extent to which a viral genotype was chimeric was quantified by the number of gene segments stemming from the parental virus that contributed fewer gene segments to the genotype. Viral genotypes were thus categorized as having between 0 and 4 minority segments. We further quantified the total number of unique viral genotypes that belonged to each of these categories. All possible genotypes in a given category were typically not observed, with the exceptions of the 0 minority segments category (which includes only the two parental genotypes) and the 1 minority segment category in mallards (**Figure 5A**). To test whether the distribution of observed genotypes across the five categories may be due to selection, we calculated the proportion of possible genotypes that were detected in each category and compared these results to a simulated dataset (**Figure 5B**). The simulated data were generated by sampling segments at random from the set of reassortant genotypes observed in each animal. Genotypes with one minority segment tended to be more prevalent in observed compared to simulated datasets, but this trend was not statistically significant owing to wide confidence intervals in the simulated data sets (**Figure 5B**). Conversely, more highly chimeric genotypes occurred less often in observed compared to simulated datasets. This trend was statistically significant for genotypes with three or four minority gene segments in guinea pigs and for those with three minority segments in mallards. Thus, observed genotype patterns likely resulted from a combination of limited availability of certain segments in a given host and purifying selection acting more strongly on reassortant genotypes in the three and four minority gene segment categories. Evidence for selection was, however, stronger in guinea pigs than mallards. In comparing mallards and guinea pigs, it is also clear that higher proportions of genotypes in each reassortant category were detected in mallards. This effect was recapitulated in the simulated dataset, however, indicating that this species-specific effect is a simple manifestation of increased genotype richness in mallards.

**Figure 5:**
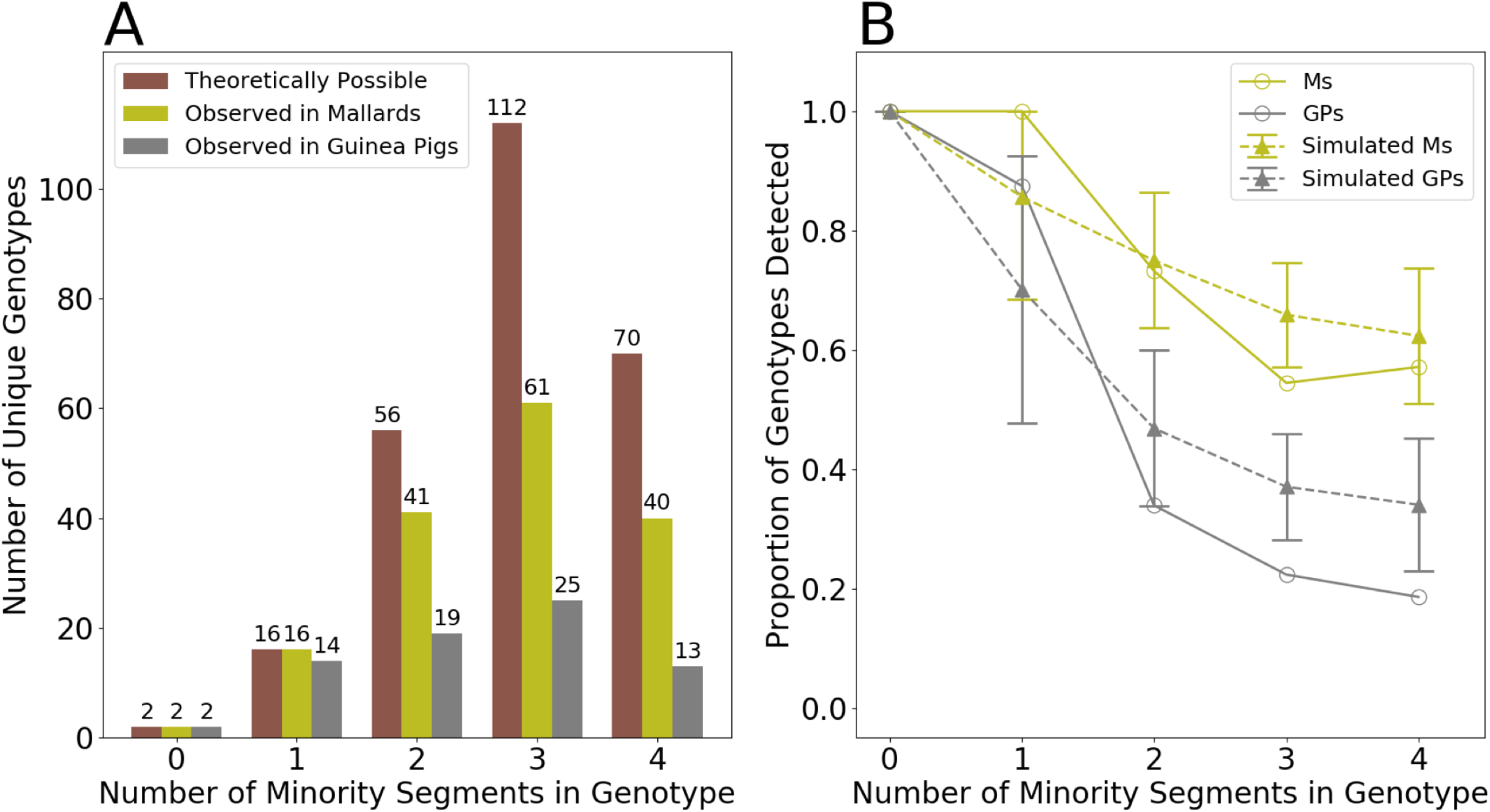
Genotype patterns detected in mallards and guinea pigs are consistent with random sampling of gene segments. A) Number of theoretically possible unique genotypes in each category (brown) and the number detected in mallards (olive) and guinea pigs (grey). Data from all time points and all individuals of a given host species are combined. Number of unique genotypes is displayed above each bar. B) Proportion of unique genotypes in each category that were detected. Observed results in mallards (solid, olive line) and guinea pigs (solid, grey line) are compared to simulated data sets generated by randomly sampling segments from reassortant viruses detected in each species (dashed lines). Error bars on the simulated data show the 95% range of calculated proportions from the 1000 simulated data sets.

## Discussion

Through experimental coinfection of mallards and guinea pigs with distinct IAVs typical of those that circulate widely in North American ducks, we examined the efficiency of IAV genetic exchange in both a natural host (mallard) and a model mammalian host (guinea pig). Robust reassortment was apparent in mallards, giving rise to highly diverse viral populations within coinfected individuals. In contrast, reassortant genotypes were less common in guinea pigs. In both hosts, temporal trends in the frequency of specific or overall reassortant genotypes were not apparent. Little recurrence of reassortant genotypes across individuals was apparent, suggesting that stochastic processes were more potent than selection in shaping within-host viral populations.

Low diversity in guinea pigs may result from relatively little opportunity for reassortment, negative selection of less fit variants, or a combination of both. Our analyses suggest that a combination of these two factors is at play in guinea pigs. If opportunity for reassortment is low, owing to low rates of coinfection, this would not only favor maintenance of parental genotypes, but would also lead to out-growth of reassortants that form and then are propagated without further coinfection. Low evenness among reassortant genotypes detected in guinea pigs is therefore consistent with limited opportunity for reassortment. Furthermore, stochastic outgrowth of reassortants would lead to over-representation of differing reassortant genotypes in unrelated individuals. In line with this expectation, recurrent detection of reassortant genotypes across different individuals was not a major feature of the dataset. One exception to this generalization was seen: a reassortant combining the NP segment of the H4N6 parent with seven segments from the H3N8 parent and was detected repeatedly in six of eight guinea pigs. More broadly, however, the reassortant genotypes detected in guinea pigs suggest that their propagation was largely a stochastic process and not the result of positive selection. Evidence of negative selection was, however, seen with the observation that highly chimeric genotypes were under represented in guinea pigs. Together, the data support roles for both stochastic and selective forces shaping reassortment patterns in guinea pigs.

In contrast to guinea pigs, the viral genotypic diversity observed in coinfected in mallards is consistent with abundant opportunity for reassortment and minimal within host selection. Several lines of evidence point to this conclusion. While parental genotypes were routinely detected, they did not predominate. Both high richness and high evenness contributed to diverse within-host viral populations. Pairwise linkage disequilibrium was low in mallards, suggesting that heterologous gene segment combinations were not strongly deleterious to viral fitness. Similarly, the reassortant gene constellations detected were largely consistent with random sampling of gene segments. The data clearly reveal mallards to be highly permissive hosts for influenza viral genetic exchange.

The evolutionary implications of robust genetic exchange within the mallard host are dependent on viral dynamics at larger scales of biological organization. For example, the extent of viral mixing within host populations is a major determinant of the frequency with which coinfections occur. Field studies indicate that this potential constraint is not strong in mallards. Within flyways, the migratory patterns of mallards and other waterfowl allow diverse IAV to be brought together at common migratory stopover points and over-wintering sites (16–19). One study revealed that, among 167 wild bird samples analyzed, 26% showed evidence of more than one subtype, indicative of mixed infection (20). Recent examination of a much broader dataset revealed mixed infection within North American waterfowl to be more common in winter, with multiple HA/NA subtypes detected in 3% of all isolates and 13% of winter isolates (16). Thus, within flyways, IAV mixing at the population level occurs readily in wild birds.

A second feature of viral dynamics that determines the potential for reassortants to impact larger scale evolutionary trends is the tightness of the bottleneck governing transmission between hosts. While IAV transmission bottlenecks are thought to be stringent in mammalian hosts (21, 22), formal analysis of the bottleneck in waterfowl has not been undertaken to date. Since IAV transmission in these birds occurs through a fecal-oral route, rather than a respiratory route as seen in mammals, the existence of a substantially wider bottleneck is plausible.

A third, and critical, factor in considering the potential for abundant within-host reassortment to impact population-level viral evolution is the distribution of fitness effects that results from reassortment. The fitness effects of both inter- and intra-subtype reassortment of human IAV is generally deleterious (8, 9). These negative fitness effects are readily explained by the disruption of inter-segment interactions: as constellations of viral genes co-evolve, the genetic underpinnings of epistatic interactions become lineage-specific and, as a result, reassortment breaks this epistasis (23). Interestingly, the consequences of genetic exchange may be very different for IAV circulating in wild waterfowl. In this ecological niche, IAV reassortment is frequently detected at the population level (20, 24–28). In particular, incongruence among the phylogenies of different IAV gene segments indicates that their evolution does not occur in concert; rather, gene constellations are routinely disrupted through reassortment. In the absence of stable gene constellations, there may be little opportunity for divergent epistatic interactions to evolve. Instead, the gene segments of avian IAV appear to exist as functionally-equivalent and readily interchangeable units (20). In stark contrast to IAV circulating in humans, reassortants that form in wild birds are not subjected to strong negative selection and circulate widely.

The observation herein of robust IAV reassortment within individual mallards strongly suggests that free-mixing and prevalent coinfection within mallard hosts contributes to the abundant IAV reassortment observed at the population level. Over multiple scales of biological organization, IAV dynamics in mallards (and likely other members of the Anseriformes) appear to overcome both spatial and selective constraints to support production, onward transmission and broad circulation of reassortant viruses.

## Materials and Methods

### Virus isolation and propagation

The viruses used in this study were originally isolated from wild bird environmental samples as part of US surveillance for avian IAV in wild birds conducted between 2006-2009 (29). Fresh wild bird feces were collected using Dacron swabs and stored in BA-1 viral transport medium. Samples were tested by RT PCR and positive samples were inoculated into the allantoic cavity of 11-day old embryonated hen eggs at 37°C for virus isolation (30). Virus subtypes were confirmed by hemagglutination inhibition and neuraminidase inhibition tests at the National Veterinary Services Laboratory (Ames, Iowa).

The H3N8 virus stock, A/wildbird/CA/187718-36/08, was propagated in hen eggs from the original environmental sample as described above. The H4N6 virus, A/mallard/CO/P66F1-5/08, was also propagated in hen eggs from the original sample but was then passaged through a mallard. A fecal sample from that mallard was propagated in hen eggs. Allantoic fluid was harvested and pooled for each virus, and aliquots were stored at −80°C prior to use. Egg Infectious Dose 50 (EID_50_) titers were determined using the Reed & Muench (1938) method.

### Whole genome sequencing and primer design

The complete viral genomes for both the H3N8 and H4N6 viruses were sequenced at the CDC Influenza Division using an Illumina sequencing platform. Briefly, viral RNA was extracted from virus stocks using the Qiagen Viral RNA Mini Kit (Qiagen) and was used as a template for multi-segment RT PCR as described previously (31). The sequence reads were assembled using the IRMA v 0.9.1 pipeline (32). The complete sequences were submitted to GenBank (MT982372-79 (H4N6); MT982380-87 (H3N8)).

Primers to distinguish H3N8 and H4N6 gene segments were designed based on these sequences (**Table 2**). The six non-HA, non-NA segments were typed using high resolution melt analysis. Primers anneal to conserved sequences of each segment and direct amplification of an approximately 100 bp region, which contains one to five nucleotide differences between the two viruses. These nucleotide differences confer distinct melting properties on the cDNA. Because the HA and NA segments are highly divergent, high resolution melt analysis is not feasible. For these segments, virus-specific primers were generated to enable genotyping by standard RT qPCR.

**Table 2:**
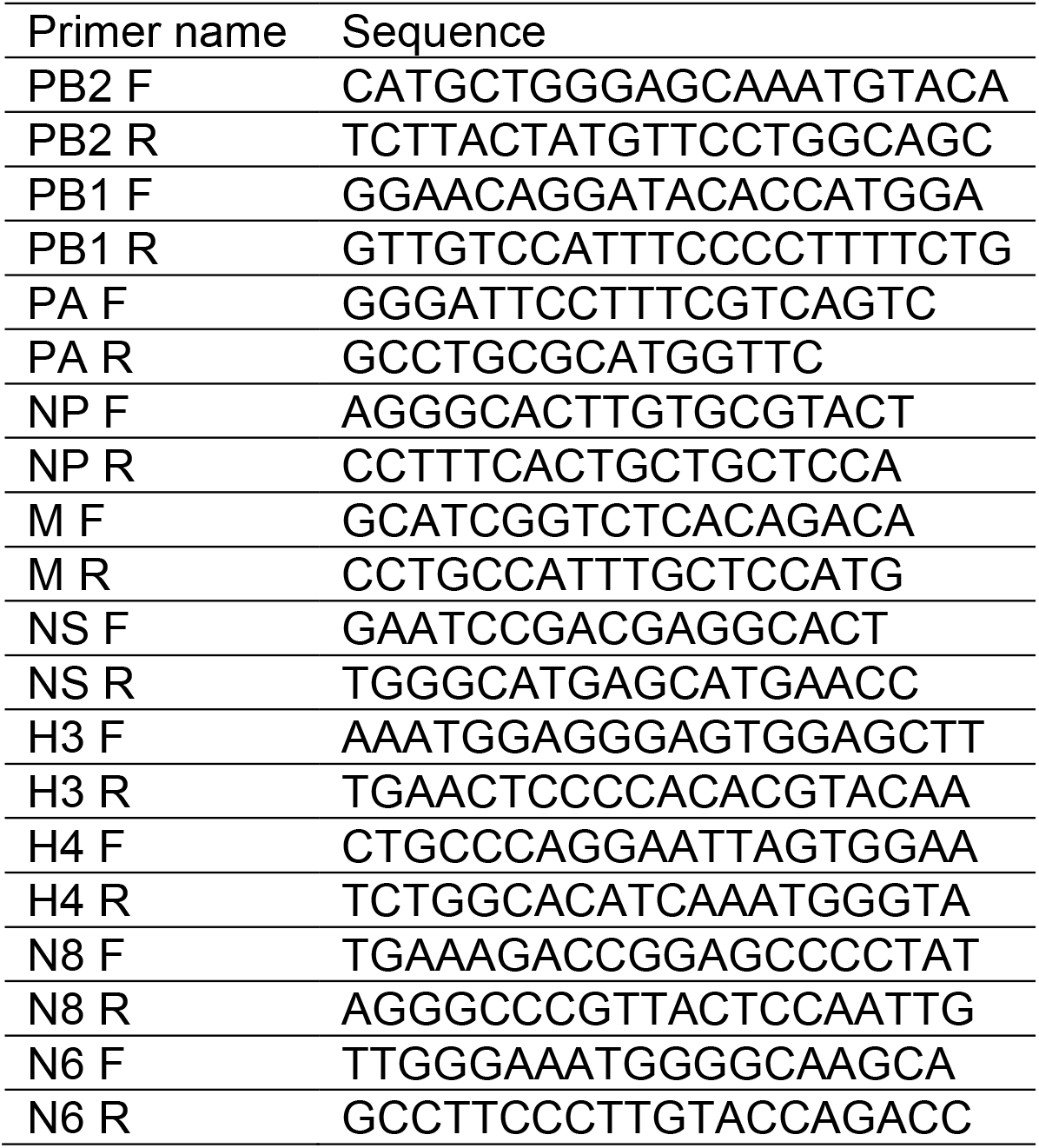
**Primers used for viral genotyping**

### Cells

Madin Darby canine kidney (MDCK) cells from Dr. Daniel Perez at University of Georgia were used for plaque assay. The cells were maintained in minimal essential medium (Gibco) supplemented with 10% fetal bovine serum (FBS; Atlanta Biologicals), penicillin (100 IU), and streptomycin (100 µg ml^−1^; PS; Corning). All cells were cultured at 37°C and 5% CO_2_ in a humidified incubator. Cells were tested monthly for mycoplasma contamination while in use.

### Animal models

All animal experiments were conducted in accordance with the Guide for the Care and Use of Laboratory Animals of the National Institutes of Health. The studies were conducted under animal biosafety level 2 containment and were approved by the Institutional Animal Care and Use Committee (IACUC) of the US Department of Agriculture, Animal and Plant Health Inspection Service, Wildlife Services, National Wildlife Research Center (NWRC), Fort Collins, CO, USA (approval NWRC QA-1621) for the mallards (*Anas platyrhynchos*) and the IACUC of Emory University (protocol PROTO201700595) for the guinea pigs (*Cavia porcellus*) studies. The animals were humanely euthanized following guidelines approved by the American Veterinary Medical Association.

### Experimental infection of mallards

Mallards were purchased from Field Trial Gamebirds in Fort Collins, CO and were approximately 5-6 months old at the time of testing. Prior to inoculation, all birds were confirmed to be negative for influenza A virus antibodies by bELISA (33) and for viral RNA by RT-PCR. During testing, all birds (n=8) were housed at the NWRC in an indoor aviary equipped with ten 2.1 m x 2.1 m x 2.4 m pens separated by 1.27 cm x 7.62 cm PVC-coated wire mesh. Birds were housed in groups of three or four per pen and could roam freely within the pen. Each pen included a food bowl, a water bowl, a small bowl of grit, and a water-filled 375 L oval stock tank for swimming and preening. The H3N8 and H4N6 inoculums were prepared to 10^5^ EID_50_/mL by diluting stocks with negative allantoic fluid. Birds were inoculated oro-choanally using a P1000 pipet with 1 ml H3N8 and 1 ml H4N6 virus preparations. Cloacal swabs were collected daily from all birds and placed in 1 ml BA-1 viral transport medium and stored at −80°C until laboratory testing.

### Experimental infection of guinea pigs

Female Hartley strain guinea pigs weighing 250–350 g were obtained from Charles River Laboratories and housed by Emory University Department of Animal Resources. Before intranasal inoculation and nasal washing, the guinea pigs were anaesthetized with 30 mg kg^−1^ ketamine and 4 mg kg^−1^ xylazine by intramuscular injection. The guinea pigs (n=8) were inoculated intranasally with the H3N8 and H4N6 virus mixture in PBS at a dose of 10^5^ pfu of each virus in a total inoculation volume of 300 µl. Animals were singly housed in filter top covered rat cages with food and water provided ad libitum. Daily nasal washes were collected in 1 ml PBS and stored at −80°C until laboratory testing.

### Determination of viral loads

Viral loads in mallard cloacal swab samples and guinea pig nasal wash samples were determined by standard plaque assay on MDCK cells. Briefly, one aliquot was thawed, mixed well and subjected to 10-fold serial dilution in PBS. Dilutions 10^−1^ to 10^−6^ were then used to inoculate confluent MDCK cells in 6-well plates. Following a one-hour attachment period, inoculum was removed, monolayers were washed with PBS and serum-free culture medium containing 0.6% Oxoid agar was overlaid onto the cells. Cultures were incubated for two days at 37°C, at which time plaques were counted and titer determined by taking into account the initial dilution of the sample. The limit of detection of the plaque assay is 50 PFU/mL, equivalent to one plaque from the 10^−1^ dilution.

### Quantification of reassortment

Reassortment levels were evaluated by genotyping 21 virus isolates per sample from mallard cloacal swabs and guinea pig nasal washes, as described previously (7). Based on positivity for infectious IAV in all eight animals, samples from days 1, 2 and 3 were chosen for evaluation of reassortment in mallards, while samples from days 2, 3 and 4 were chosen for guinea pigs.

Briefly, plaque assays were performed on MDCK cells in 10 cm diameter dishes to isolate virus clones. Serological pipettes (1 ml) were used to collect agar plugs from well-separated plaques into 160 µl PBS. Using a ZR-96 viral RNA kit (Zymo), RNA was extracted from the agar plugs and eluted in 40 µl nuclease-free water (Invitrogen). Reverse transcription was performed with a universal IAV primer (31) using Maxima reverse transcriptase (RT; Thermofisher) according to the manufacturer’s protocol. The resulting cDNA was diluted 1:4 in nuclease-free water, each cDNA was combined with segment-specific primers (**Table 2**) and Precision Melt Supermix (Bio-Rad) and analyzed by qPCR using a CFX384 Touch real-time PCR detection system (Bio-Rad). The qPCR was followed by high-resolution melt analysis to differentiate the H3N8 and H4N6 amplicons of the six non-HA, non-NA segments. Precision Melt Analysis software (Bio-Rad) was used to determine the parental virus origin of each gene segment based on the melting properties of the cDNA fragments and comparison to H3N8 and H4N6 virus controls. For the HA and NA gene segments, Ct values were used to determine the origin of each segment. Ct values >= 32 were considered negative. Each plaque was assigned a genotype based on the combination of H3N8 and H4N6 genome segments, with two variants of each of the eight segments allowing for 256 potential genotypes. In a minority of samples, the origin of one or more gene segments could not be determined and the genotype from that clonal isolate was removed from the analysis.

### Software

All calculations, plotting, and machine learning was done through Python 3 (34). Packages used included matplotlib (35), NumPy (36), pandas (37), SciPy (38) and statsmodels (39).

### Genotype Frequencies, Richness, Diversity, and Evenness

Here, a virus genotype is defined as a unique combination of the eight IAV segments, where each segment is derived from either the H3N8 or H4N6 parental virus. In total, there are 2^8^ (=256) possible unique genotypes. Genotype frequencies were calculated for each sample by dividing the number of appearances of each genotype by the total number of clonal isolates available for that sample.

Genotype richness (*S*) is given by the number of unique virus genotypes in a sample. Genotype diversity was quantified using the Shannon-Wiener Index:

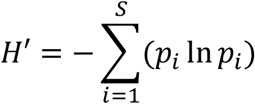

where *S* is genotype richnessand *p*_*i*_ is the frequency of genotype *i* in the sample (40). With a range between 0 and 1, genotype evenness is defined as the extent to which genotypes are evenly present in a sample, with 1 corresponding to maximum evenness (41). Evenness is given by:

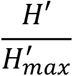

where maximum genotype diversity 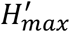 evaluates to In *S* in a sample with genotype richness *S*. Evenness values were calculated both with and without the parental genotypes included in the samples. In the evenness calculations that excluded parental genotypes, genotype frequencies, richness *S*, diversity *H*′ and maximum diversity 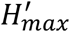 values were all recalculated using the subset of clonal isolates that were not parental genotypes.

### Linkage Disequilibrium

Linkage disequilibrium *D* is a population genetic measure that quantifies the extent of non-random association of alleles at two or more loci in a population (42). Here we use *D* to characterize non-random associations of gene segments. *D* was calculated for all 8-choose-2 (= 28) possible two gene segment combinations using the formula:

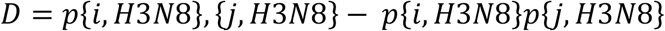

where *p*{*i, H*3*N*8} is the proportion of clonal isolates that have gene segment *i* derived from parental virus H3N8 *p*{*j, H*3*N*8} is the proportion of clonal isolates that havegene segment *j* derived from parental virus H3N8, and *p*{*i, H*3*N*8}, {*j, H*3*N*8} is the proportion of the clonal isolates have both gene segments *i* and *j* derived from parental virus H3N8. Calculations of *D* using H4N6 as the reference virus instead of H3N8 yield equivalent results.

### Minority Segments, Unique Genotypes, and Simulation

To identify potential purifying selection in mallards and guinea pigs, an analysis of the genotypes in terms of the number of minority segments was conducted. We define a minority segment as a segment derived from the virus that contributes fewer gene segments to a genotype. Each observed genotype can thus be classified as falling into a category of containing between 0 and 4 minority gene segments. Figure 5A shows the number of virus genotypes that fall into each of these minority gene segment categories.

To determine if selection acted against more chimeric genotypes, the proportion of genotypes in each minority gene segment category was first calculated by dividing the observed number of genotypes in that category by the number of theoretically possible genotypes in that category (Figure 5B). To determine whether these calculated proportions deviated from what would be expected by chance, we generated simulated data for both mallards and guinea pigs that reflected chance expectations. Specifically, during a round of simulation, all clonal isolates containing parental genotypes were first excluded from each of the samples. The remaining clonal isolates were then randomly permuted by gene segment to generate a simulated data set. If the simulated data set contained any parental genotypes, the data set would be tossed and another one generated. 1000 simulated data sets were generated in total. Each of these resultant simulated data sets were analyzed similarly to the observed data sets.

### Statistical Measures

All confidence intervals were calculated via the proportion_confint method from the statsmodels package (39). Unpaired t-tests were conducted via the ttest_rel and the ttest_ind methods respectively from the SciPy package (38). Mann-Whitney U tests were conducted with the mannwhitneyu method and ANOVA tests were conducted with the f_oneway method, both of which are found in the SciPy package (38).

## Acknowledgements

This work was funded by the National Institute of Allergy and Infectious Disease through R01 AI127799 and the Centers of Excellence for Influenza Research and Surveillance (CEIRS) contract no. HHSN272201400004C.

## Supplementary Figure Legends

**Supplementary Figure 1: Viral genotypes detected in mallards and guinea pigs**.

Viral genotypes detected in mallards (A) and guinea pigs (B) are depicted with H4N6 parental origin in blue and H3N8 parental origin in orange. Each table corresponds to a sample collected from an individual animal on a given day, with columns representing the eight viral gene segments ordered 1-8 from left to right and rows showing clonal isolates. Tables are sorted such that any parental genotypes appear at the top (H4N6) or bottom (H3N8).

**Supplementary Figure 2: Viral genotype frequencies detected in mallards and guinea pigs**. Stacked plots showing frequencies of unique genotypes detected in each animal over time. The lowermost two sections, in blue and orange, represent the H3N8 and H4N6 parental genotypes, respectively.

