## Supplementary figures and images for "Avian influenza A viruses reassort and diversify differently in mallards and mammals"

### Supplemental Figures

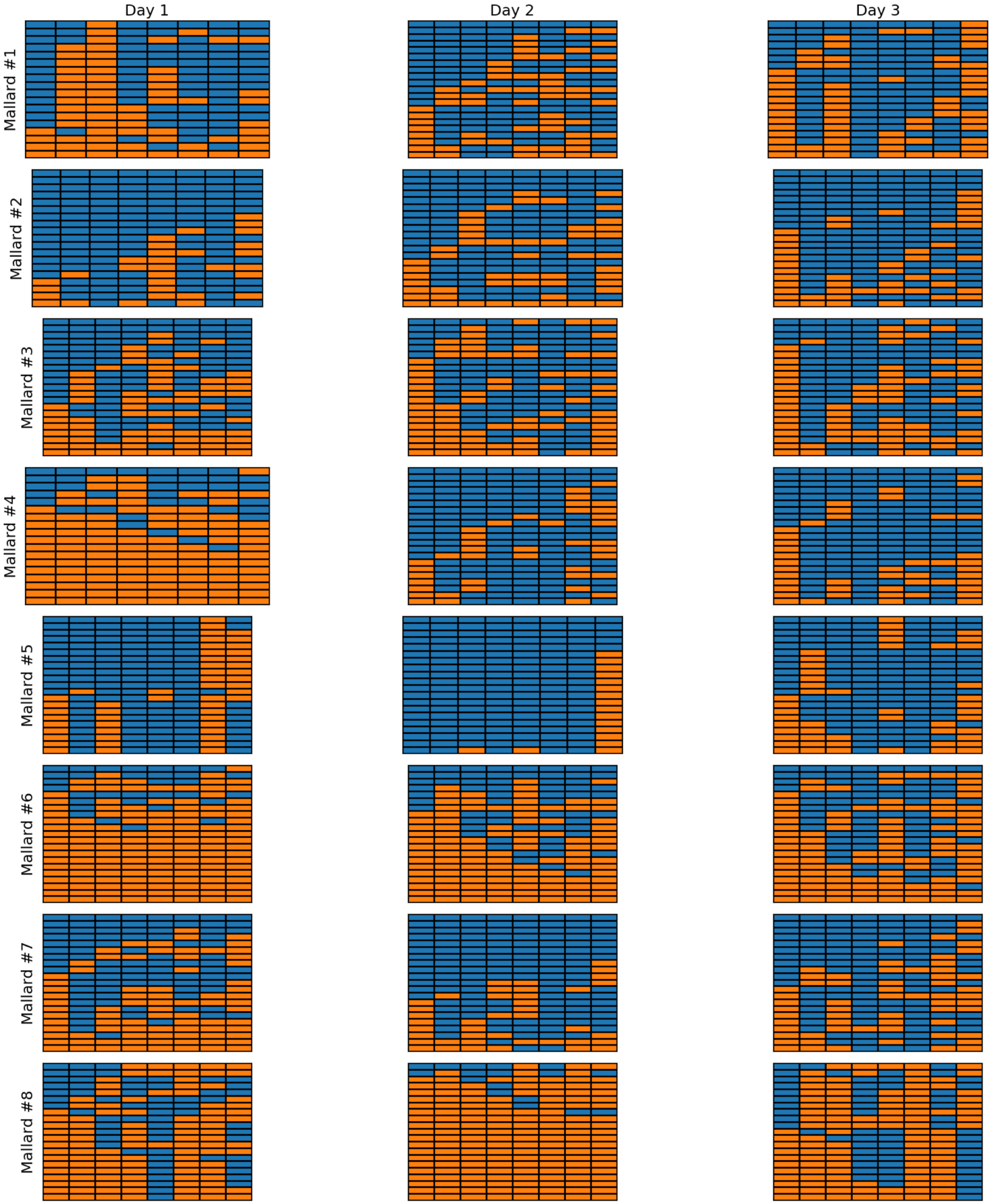

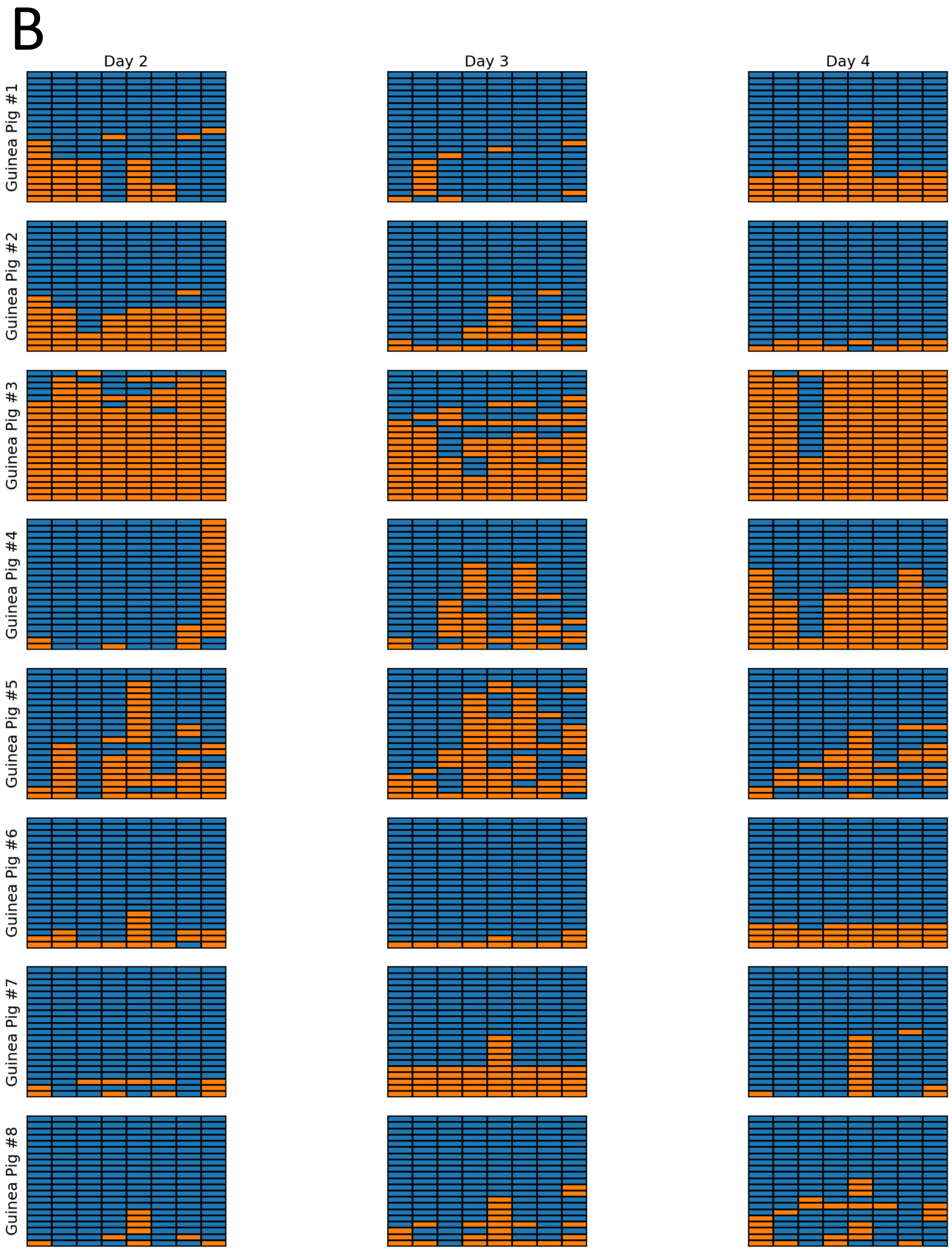

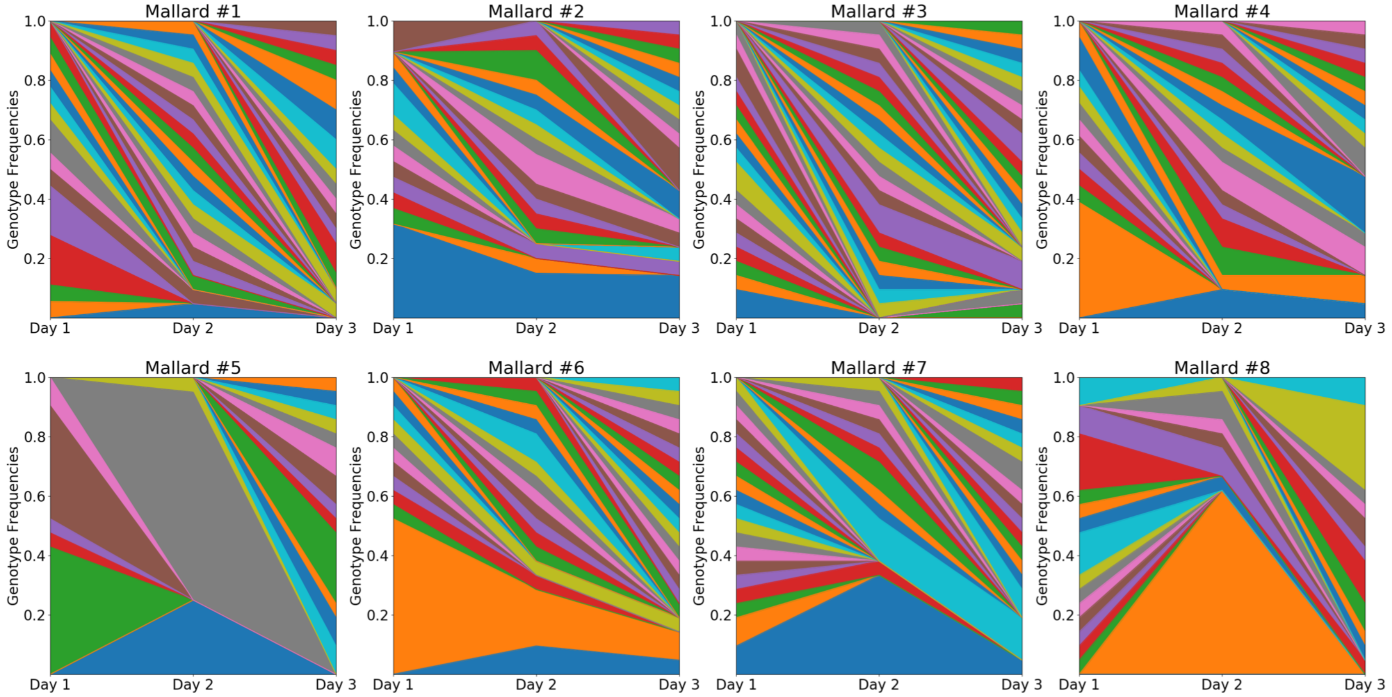

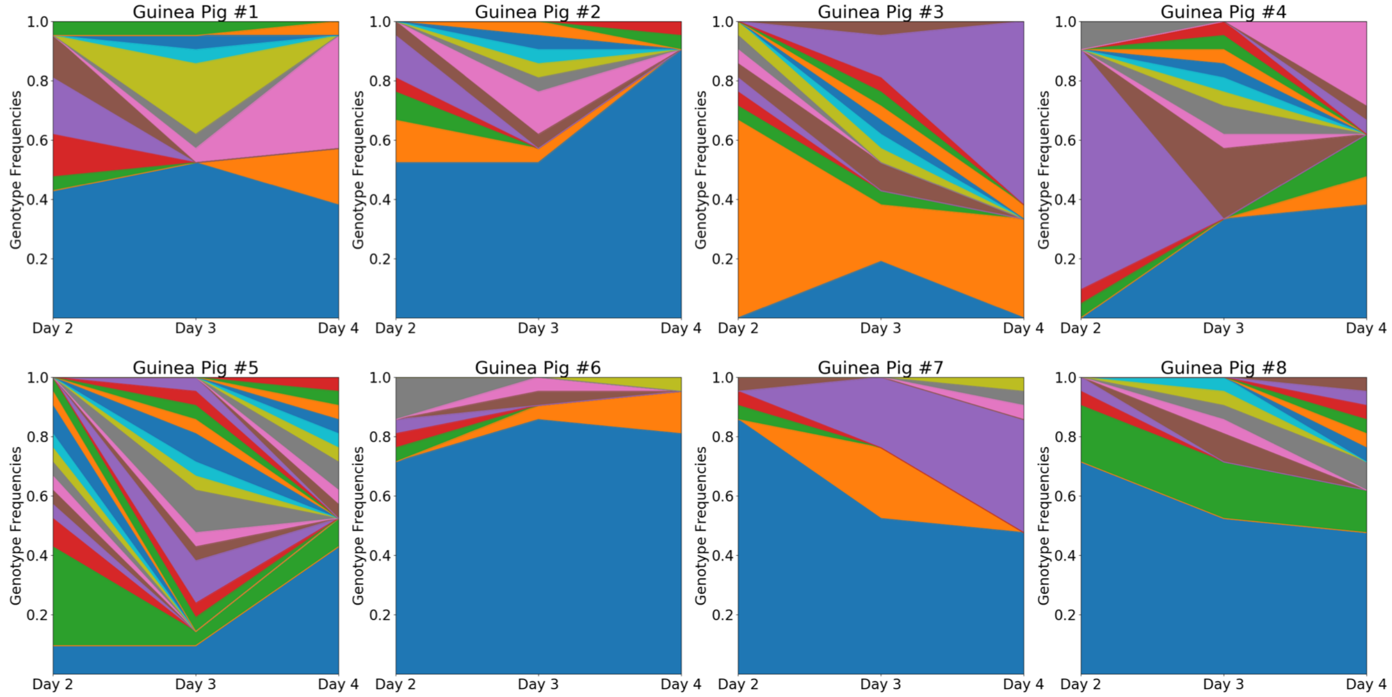
